# RapidMACS: MACS3-identical peak calling, 50× faster

**DOI:** 10.64898/2026.08.13.744681

**Authors:** Ling-Hong Hung, Ka Yee Yeung

## Abstract

**Motivation:** MACS3 is a comprehensive peak-calling toolkit whose subcommands span single- and paired-end data, narrow and broad peaks, and a range of signal-track utilities. However, many ATAC-seq and multiomic pipelines, including our own, use just one of those capabilities: narrow peak calling. A single analysis may call peaks under different conditions, so a per-call saving is multiplied and the time recovered can be substantial. Furthermore, we wanted an efficient and embeddable narrow peak caller that could be integrated directly with our Chromap Suite aligner, so we wrote RapidMACS.

**Results:** RapidMACS is a narrow peak caller optimized for this purpose, building its signal tracks in a single lazy sweep that avoids a global sort, processing chromosomes in parallel, and keeping intermediates in memory rather than creating temporary files. In benchmarks involving single-cell ATAC-seq, bulk ATAC-seq, ChIP-seq, and CUT&RUN, it is 3.5–51× faster than MACS3 v3.0.3 depending on the applications and number of threads used. More importantly, the output is byte-identical. This means that any downstream analysis using RapidMACS will produce identical results to those using MACS3. The speed gains are largely due to these algorithmic changes rather than the choice of language, because MACS3’s peak-calling code is itself compiled (Cython). RapidMACS depends only on htslib and zlib and links as a small static archive without a Python or Cython runtime.

**Availability and implementation:** RapidMACS is open source under the MIT license at https://github.com/morphic-bio/rapidmacs; the results reported here are from v1.0.1. It provides both a standalone CLI executable (`rapidmacs`) and a linkable C++ library. Prebuilt containers for x86-64 and arm64 are published as `biodepot/rapidmacs` on Docker Hub.

**Supplementary information:** Supplementary data are available at Bioinformatics online.

## 1. Introduction

MACS (Zhang et al., 2008) has been the standard narrow peak caller for chromatin assays for nearly two decades, and it has remained so even as other platforms began including their own. For example, Cell Ranger ARC’s ATAC peaks are wider, and downstream analyses routinely re-call narrow peaks with MACS through Signac (Stuart et al., 2021), ArchR (Granja et al., 2021), and SnapATAC2 (Zhang et al., 2024); ENCODE’s ATAC-seq and transcription-factor peak sets are MACS calls (ENCODE Project Consortium, 2012). Narrow peaks are necessary for these analyses: accessibility and sequence-specific factor binding produce localized signals a few hundred bases wide. Narrow peak callers such as MACS3 return the summit, the single base of highest pileup within a peak, which is used to define a peak region. Motif enrichment and footprinting are anchored on summits—ArchR, for example, extends each summit by ±250 bp into the 501 bp fixed-width, non-overlapping peak set to create its cell-by-peak count matrix. Multiple MACS3 calls are often required. One cell contributes at most a fragment or two at any locus, so peaks can be called only after cells are aggregated to form a pileup. These pileups cannot be recovered from an earlier call’s output, so every grouping means another MACS invocation: ArchR and SCENIC+ both call it once per group and merge the results (Bravo González-Blas et al., 2023).

### Our contributions

Our work in the MorPhiC consortium (Adli et al., 2025) required an efficient open-source peak caller that produced narrow peaks needed for downstream analyses. MACS3 is open source (BSD-3-Clause) and already fast, being implemented in compiled Cython against the CPython runtime. However, we had to invoke it separately, which meant writing intermediate files and reading them back. Linking MACS3 directly was not a feasible solution because Cython requires the Python interpreter. The alternative of invoking `macs3` as a subprocess would not have solved the problem of writing intermediate files. We therefore wrote RapidMACS so that it could be linked directly into our Chromap Suite aligner, which extends Chromap (Zhang et al., 2021), and into the rest of our pipeline for ATAC-seq and multiomic analyses. Although we only needed to call narrow peaks in paired-end fragments from ATAC-seq, both ChIP-seq and CUT&RUN use essentially the same methodology, so we also added support for them. In addition to the linkable C++ library that we needed, we also added a CLI so that RapidMACS can also be used as a standalone program in any workflow that currently invokes `macs3 callpeak` to call narrow peaks.

## 2. Implementation

### Scope and limitations

RapidMACS calls narrow peaks, with or without a matched input or IgG control, from fragment, BAMPE, BEDPE, or single-end BED input. It does not support any other MACS3 subcommands; broad calling in particular is a separate algorithm, with its own pair of cutoffs and region-linking pass. Single-end input additionally requires `–nomodel`, since MACS3’s fragment-size model building is not implemented. This focus allowed us to optimize RapidMACS extensively; the changes are described next.

### Methodological changes

MACS3’s peak-calling path is compiled Cython rather than interpreted Python and the speed gains are not primarily due to the language differences but to three methodological changes. Two are structural: first, chromosomes are independent once the fragments are grouped, so RapidMACS calls them concurrently rather than in sequence, and second, it holds each chromosome’s tracks in memory where MACS3 serializes them to a temporary file between passes. The third is how the pileups themselves are built, which is the central algorithm and the subject of the rest of this section.

To call a peak we need two tracks: the treatment track, holding the number of fragments covering each base, and the local background it is compared against. Both RapidMACS and MACS3 store these tracks as runs of constant value rather than as per-base arrays—what differs between the implementations is how those runs are built.

For single-cell ATAC-seq, the input is a fragment TSV file, from Cell Ranger or from Chromap. These are sorted by start position, but not by end position. MACS3 sorts the ends, then walks the sorted start and end arrays together to build the treatment track. It builds the background track the same way, from endpoints widened into a 10 kb window, which yields the average coverage over that window, with the genome-wide mean as a floor.

RapidMACS replaces that global sort with a lazy one that orders only the ends of the fragments spanning the current position. It keeps those pending ends in a heap and pops them as the sweep reaches them, which yields the same sequence of events a full sort would. A single sweep generates both the treatment and the background track, using a separate heap for each. This trades *O*(*n* log *n*) for *O*(*n* log *k*), and on the single-cell ATAC data the two are far apart: *n* is 53.0 M fragments, while *k*—the number spanning a given position—averages 4 for the treatment track and 510 for the 10 kb background window. Each heap is therefore a few kilobytes and stays in first-level cache for the whole sweep.

For bulk ATAC-seq, ChIP-seq, and CUT&RUN, coordinate-sorted BAMPE input is the norm, and sorting on the start coordinate very nearly sorts the ends along with it: both endpoints come from the same read pair, and the fragment between them is only a few hundred bases. Sorting an array already that close to order is cheap, so there is little for a lazy sweep to save. The heap method remains available through flags, but our preliminary tests showed it to be no faster, and slightly slower on ChIP-seq, because of the overhead of creating and managing the heap. RapidMACS therefore uses a single global sort on BAMPE by default. In all cases the two tracks are then overlaid, and each run over which both are constant is scored once by the Poisson tail probability of seeing that many fragments under the local background. A peak is then a stretch of runs clearing the threshold that is uninterrupted by a gap wider than a set width, and kept only if what results exceeds a minimum length.

### Two ways to use it

As a library, RapidMACS builds to a static archive depending only on htslib (Bonfield et al., 2021) and zlib, with no Python or Cython runtime. Its entry point, `RunMacs3FragPeakPipelineFrom` SortedIterator, asks the caller for fragments one at a time, so they can come from a file or straight out of an array the host is already holding. As a standalone program, `rapidmacs` reads a Chromap/ARC fragments TSV, a single-end BED, or, via htslib, BAMPE or BEDPE, and writes narrowPeak and summit files in place of `macs3 callpeak`.

### Open source made exact parity possible

We wrote RapidMACS from MACS3’s published description rather than porting its code. No MACS3 code is incorporated and RapidMACS is independent of MACS3’s codebase. At every stage the development procedure was the same: run the new implementation and the reference MACS3 implementation on the same input and compare the output files. RapidMACS was written with AI-assisted programming under human direction, following the approach we describe for STAR Suite (Hung and Yeung, 2026b); the stage-by-stage output comparison against a reference implementation also verifies work produced by the AI agents.

MACS3’s documentation leaves two things unclear: the tie-break it applies when several positions inside a peak share the maximum pileup, and the 32-bit arithmetic of its score tables. Both must be matched exactly for the output files to agree. Because MACS3 is open source we could read and instrument v3.0.3 to establish what it does in each case. Against a closed peak caller we could still have checked byte-identity, since that needs only the two output files. What we could not have determined was why they differed, and the likely outcome would have been a tool that agreed closely but not exactly.

## 3. Results

We validated on four full public datasets covering single-cell ATAC-seq, bulk ATAC-seq, ChIP-seq, and CUT&RUN: the 10x 3K PBMC Multiome ATAC channel (10x Genomics, 2021); ENCODE bulk ATAC-seq in GM12878 (ENCSR095QNB) (ENCODE Project Consortium, 2020); ENCODE CTCF ChIP-seq in NCI-H929 with its matched input control (ENCFF686KKV/ ENCFF181CXT) (ENCODE Project Consortium, 2021); and K562 CTCF CUT&RUN with matched IgG (SRR14255054/ SRR14255053) (Ng et al., 2023). The public sets were ingested directly from the fragments or sorted BAMPE files provided. As a further control, the two ATAC datasets were additionally realigned from the same reads with Chromap Suite, because that is how ATAC data reaches the peak caller in our MorPhiC workflows. Each ATAC dataset therefore appears twice in Table 1—the same reads, aligned once by its source and once by Chromap—which with ChIP-seq and CUT&RUN makes six configurations.

**Table 1.** Complete-program wall time against MACS3 v3.0.3 across six input configurations. Each row is one benchmark input, used either as its source distributes it or realigned by Chromap Suite from the same FASTQ files. Times are wall-clock seconds for the complete command-line program, medians of three recorded warm-cache runs after one unrecorded warm-up. All runs ran on a single server (i9-13900KF, 128 GB RAM) with every input, output, and temporary file on the same NVMe device. *T* is the number of RapidMACS threads. MACS3’s peak-calling path is single-threaded so each row has only one MACS3 time. Parenthesized values are speedups against it. Peak counts differ between rows because the inputs differ. Within every row, and in every run, both programs produced byte-identical narrowPeak and summit files. Stage specific timings are in Supplementary Section S5.

| Assay and input | MACS3 | RapidMACS |  |  | Peaks |
| --- | --- | --- | --- | --- | --- |
| | v3.0.3 | $T=1$ | $T=4$ | $T=24$ | |
| scATAC-seq, 10x fragments | 747.55 | 58.93 (12.7×) | 24.44 (30.6×) | 16.35 (45.7×) | 50,003 |
| scATAC-seq, Chromap fragments | 758.62 | 59.81 (12.7×) | 25.58 (29.7×) | 16.31 (46.5×) | 50,274 |
| Bulk ATAC-seq, ENCODE BAM | 146.63 | 41.94 (3.5×) | 13.21 (11.1×) | 7.47 (19.6×) | 74,000 |
| Bulk ATAC-seq, Chromap BAM | 122.92 | 32.51 (3.8×) | 10.84 (11.3×) | 6.56 (18.7×) | 78,574 |
| ChIP-seq, ENCODE BAM + input control | 616.90 | 97.08 (6.4×) | 27.38 (22.5×) | 12.18 (50.6×) | 33,985 |
| CUT&RUN, GEO BAM + IgG control | 64.74 | 9.94 (6.5×) | 2.99 (21.7×) | 2.01 (32.2×) | 9,097 |

On all six, every narrowPeak and summit field is byte-identical to MACS3 v3.0.3—intervals, peak names, summit offsets and coordinates, signal values, *p*-scores and *q*-scores—and the full-file MD5 sums match. By identical, we mean that literally: the same bytes, not the same peaks within a tolerance. Anything computed from a RapidMACS peak or summit file—peak annotation, differential accessibility, motif enrichment, a cell-by-peak count matrix—returns exactly what it would have returned from the MACS3 file, because it is handed the same input file.

Table 1 gives the complete running time for every configuration. RapidMACS is 3.5–51× faster depending on assay, input, and thread count, and all 72 measured runs reproduced the byte-identical output described above. Single-cell ATAC-seq has a greater performance gain because it takes fragment inputs and can use the lazy heap based sweep to avoid an expensive global sort (12.7× faster on a single thread). The other assays take sorted BAMPE as input. Without the advantage of avoiding an expensive global sort, RapidMACS is only 3.5–6.5× faster on a single thread. Chromosome-level parallelism then multiplies both, by a further 3.6–8.0× at 24 threads. The BAMPE configurations scale better with threads because a BAM file is a series of independently inflatable BGZF blocks, so decompression itself parallelises. A plain gzip fragment file is one deflate stream whose decoder state carries from the first byte to the last, leaving no offset a second thread could start from, and RapidMACS inflates it serially. The higher throughput does come with a modest increase in memory usage. Peak resident set size is 0.46–5.40 GiB against 0.35–2.74 GiB for MACS3, roughly 1.2–2.5 times as much and at most about 2.7 GiB more.

One property of the released software affects the two assays with controls. MACS3 v3.0.3 writes each chromosome’s control pileup to a fixed file name before returning it. As far as we can tell, nothing reads the file back. We found no option to disable it, so we built a version without those four lines. For ChIP-seq, whose control is the larger of the two at 39.3 M fragments, the writing takes 207.6 s, a third of MACS3’s 616.90 s run. Both builds call identical peaks; only the writing differs. Against the build without it RapidMACS is 33.6× faster at 24 threads rather than 50.6×, which leaves single-cell ATAC-seq the fastest configuration at 46.5×. Supplementary Section S4 gives both in full. Treatment-only runs never reach that code. The complete 72-run matrix and the per-stage timings are given in Supplementary Sections S2–S5.

## 4. Applications

RapidMACS is provided as a libRapidMACS library as well as a standalone program because peak calling sits in the middle of pipelines rather than at the end of them. A pipeline that maps reads and then calls peaks by invoking `macs3` has to write its fragments to disk, start a Python process, and have that process read them back. For the single-cell ATAC dataset used here this involves transferring 2.3 GB of data going back and forth between processes via writing and reading temporary files. Linking to the library removes this overhead: an aligner then hands libRapidMACS the fragments it has just produced, in the same process directly through its C++ API. That is how we use it. Chromap Suite links RapidMACS to call ATAC peaks during the mapping pass, and Multiomics Suite composes that with transcriptomic processing to run alignment, peak calling, and cell calling for joint RNA and ATAC in one invocation of one binary (Hung and Yeung, 2026a).

The standalone program serves the more common use case: re-calling peaks that a pipeline has already produced. We verified RapidMACS against the peak-calling commands issued by the three tools that do it most: Signac, which calls MACS to replace Cell Ranger ARC’s wider peaks on multiome data (Stuart et al., 2021); ArchR, which calls it once per pseudobulk replicate (Granja et al., 2021); and SCENIC+, which assembles a consensus enhancer set from per-cluster pseudobulk (Bravo González-Blas et al., 2023). All three use `macs2` by default but can use `macs3` if desired. We tested RapidMACS against the MACS3 v3.0.3 commands and confirmed that the narrowPeak and summit files are byte-identical.

## 5. Acknowledgments

We thank X. Shirley Liu’s group, where MACS was originally developed, and Tao Liu and the MACS3 contributors, who have maintained and extended it since, for sustaining MACS over many years and for releasing it under a permissive open-source license: being able to read and run the reference implementation is what made byte-level parity possible. We also thank the htslib authors.

## 8. Data availability

Source, build instructions, and the parity harness are at https://github.com/morphic-bio/rapidmacs. All benchmark inputs are public; accessions and checksums are listed in Supplementary Section S1, and the complete 72-run benchmark matrix is given in Supplementary Section S3. The audit reproducing the Signac, ArchR, and SCENIC+ invocations, with the pinned upstream revisions and per-file checksums, is at `paper/WORKFLOW-COMPATIBILITY.md`.

## Supplementary Material

### S1 Test datasets and output identity

**Table S1.**
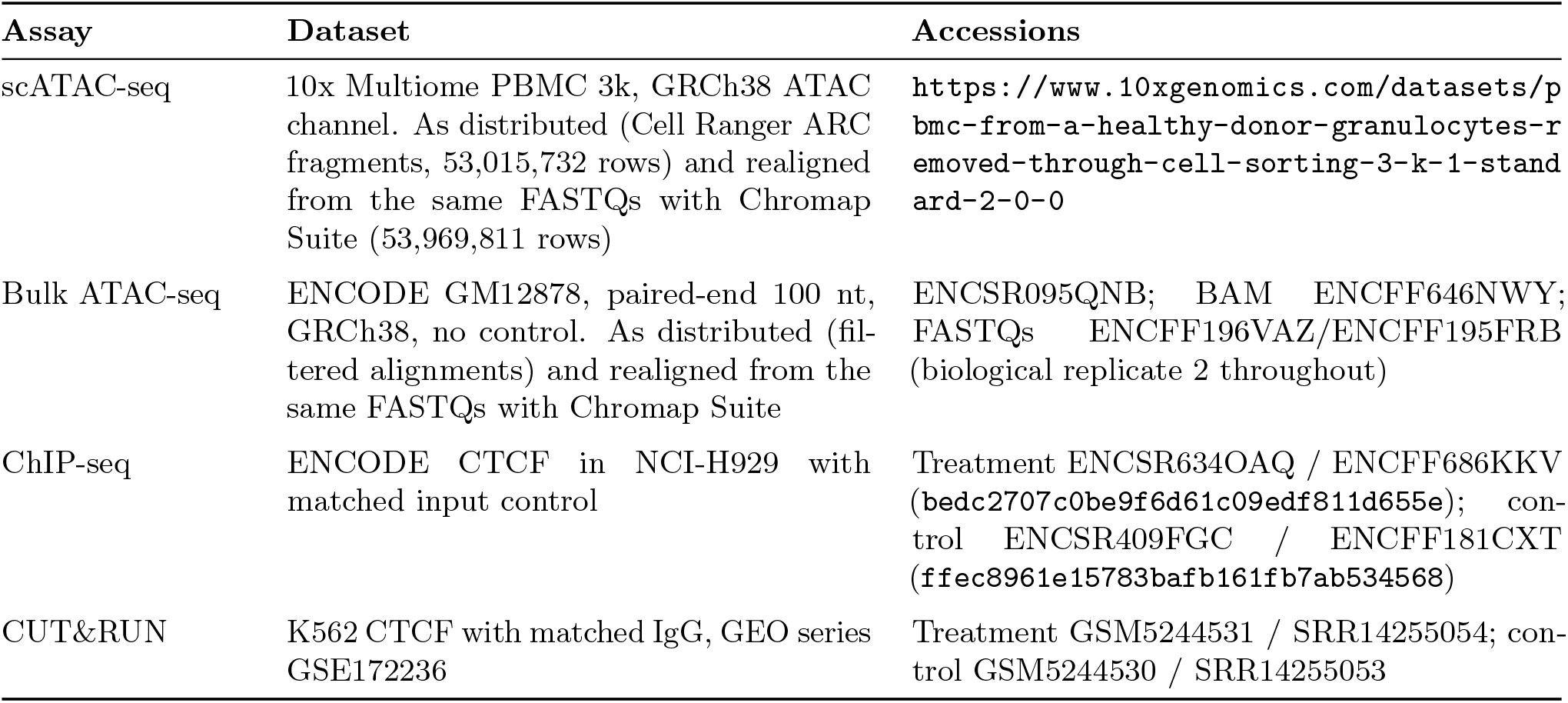
Benchmark datasets. All inputs are public. Each dataset is used in the form its source distributes it; the two ATAC datasets are additionally realigned from the same reads with Chromap Suite, giving six configurations in total. Within every configuration both tools consume the identical file, and alignment is excluded from all timings.

#### Output identity

Both programs write the same files. The MD5 sums below are of the complete narrowPeak and summits files, not of any subset of their columns, and are identical for MACS3 v3.0.3 and RapidMACS on every one of the 72 measured runs. The benchmark harness aborts if they ever differ, so no timing in this paper can come from a build that does not reproduce the output.

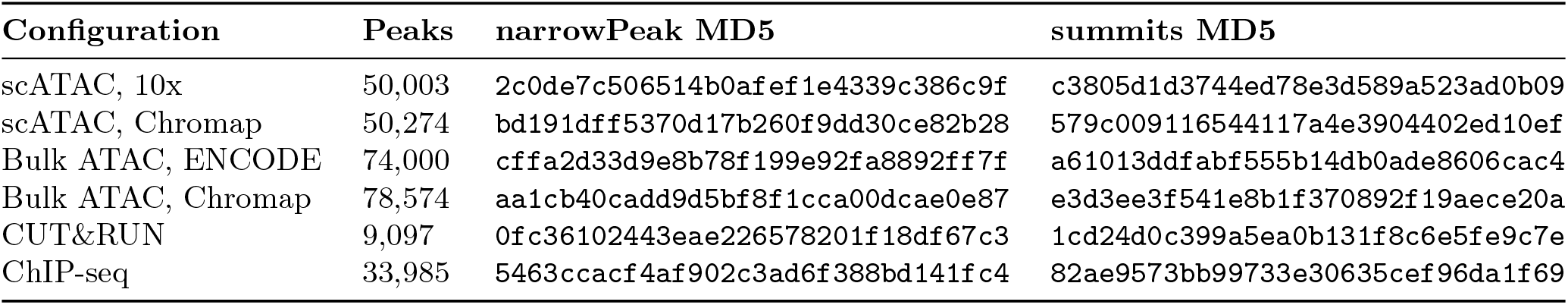

#### Compared commands

~~~
# ATAC
macs3 callpeak -t fragments.tsv.gz -f FRAG -g hs -n atac_full \
 -p 1e-5 --min-length 200 --max-gap 30 --outdir OUT
rapidmacs --preset shiftTAG -f FRAG -t fragments.tsv.gz -g hs \
 -n atac_full --peak-caller-threads T -p 1e-5 \
 --bdgpeakcall-min-len 200 --bdgpeakcall-max-gap 30 \
 --macs3-frag-narrowpeak OUT/atac.narrowPeak \
 --macs3-frag-summits OUT/atac.summits.bed
# ChIP-seq / CUT&RUN (matched control)
macs3 callpeak -t treat.bam -c control.bam -f BAMPE -g hs -n NAME \
 -q 0.05 --outdir OUT
rapidmacs --preset chip -t treat.bam -c control.bam -g hs -n NAME \
 --peak-caller-threads T --out-prefix OUT/NAME
# Bulk ATAC-seq (treatment only)
macs3 callpeak -t treat.bam -f BAMPE -g hs -n NAME -q 0.05 --outdir OUT
rapidmacs --preset cutrun -f BAMPE -t treat.bam -g hs -n NAME \
  --peak-caller-threads T --out-prefix OUT/NAME
~~~

### S2 Provenance of the measurements

The benchmark unit is the standalone executable, including input parsing, peak calling, and narrow-Peak/summits output; internal library-only timings are diagnostic and are not the reported comparison. For every assay/tool/thread configuration the harness runs one unrecorded warm-up followed by three recorded repetitions, and reports the median wall time and median maximum RSS with the observed range. Wall time and maximum RSS come from GNU `/usr/bin/time`; each run writes to a separate directory.

Warm-cache measurement is deliberate. Dropping the Linux page cache requires privileged, host-wide mutation and would disturb unrelated jobs on the shared benchmark host; a full warm-up per configuration gives both implementations the same cache state and avoids charging cold-cache variance to the peak caller. Stage profiling is enabled consistently for RapidMACS runs and its small overhead is included in the reported RapidMACS wall times. All measurements come from one 13th-generation Intel Core i9-13900KF host with no competing CPU- or I/O-intensive jobs. Every input, output directory, and temporary file resided on the same NVMe solid-state device, so neither program was advantaged by storage placement.

### Which RapidMACS and MACS3 builds were measured

Every RapidMACS timing reported in this paper is of v1.0.1. Results describes the extra file that MACS3 v3.0.3 writes on every run that has a control. ChIP-seq, CUT&RUN, and scATAC 10x were measured against the released build. The other three configurations are treatment-only, never reach that code, and were measured against a build with those four lines removed, so the executed path is the same either way. Running scATAC 10x against both bounds the residual difference at 0.13%: 747.55 s released against 748.49 s.

### S3 Full benchmark matrix

### S4 MACS3 with the extra file removed

Table S3 repeats the two controlled measurements against a MACS3 build with the four lines described in Results removed. The patch is checked in at `paper/macs3-3.0.3-no-control-dump.patch`. Both builds produce identical narrowPeak and summits files, so the difference is entirely the cost of writing the file. It is much the larger on ChIP-seq, whose 39.3 M-fragment control dwarfs CUT&RUN’s 2.4 M: 207.64 s, a third of the released build’s run. Removing it leaves single-cell ATAC-seq the fastest configuration in the paper, at 46.51×.

**Table S2.**
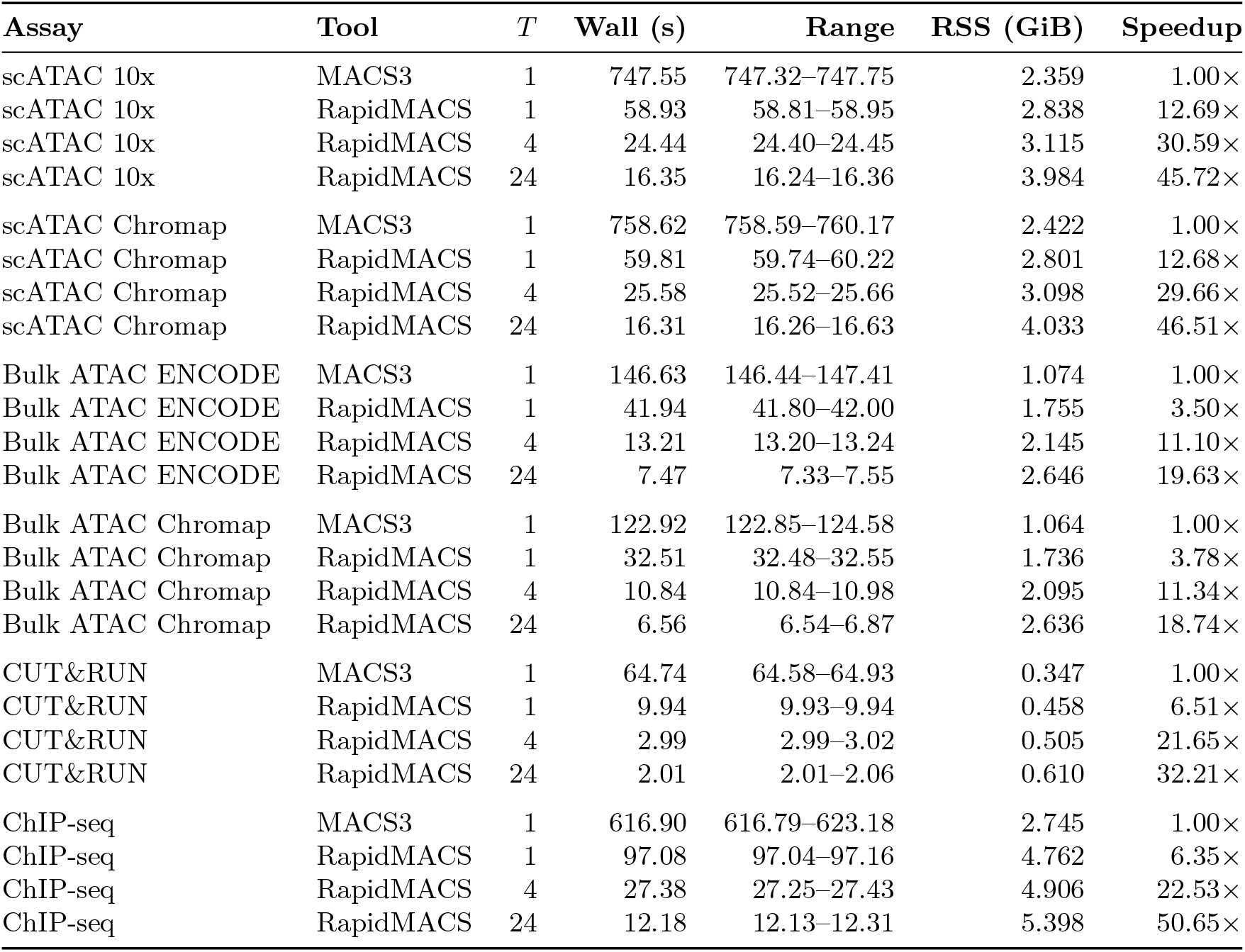
Complete-CLI medians with observed ranges over three warm-cache runs after one unrecorded warm-up. Every configuration produced narrowPeak and summit files byte-identical to MACS3 v3.0.3. Speedups are against that configuration’s own MACS3 time.

| Assay | Tool | $T$ | Wall (s) | Range | RSS (GiB) | Speedup |
| --- | --- | --- | --- | --- | --- | --- |
| scATAC 10x | MACS3 | 1 | 747.55 | 747.32–747.75 | 2.359 | 1.00× |
| scATAC 10x | RapidMACS | 1 | 58.93 | 58.81–58.95 | 2.838 | 12.69× |
| scATAC 10x | RapidMACS | 4 | 24.44 | 24.40–24.45 | 3.115 | 30.59× |
| scATAC 10x | RapidMACS | 24 | 16.35 | 16.24–16.36 | 3.984 | 45.72× |
| scATAC Chromap | MACS3 | 1 | 758.62 | 758.59–760.17 | 2.422 | 1.00× |
| scATAC Chromap | RapidMACS | 1 | 59.81 | 59.74–60.22 | 2.801 | 12.68× |
| scATAC Chromap | RapidMACS | 4 | 25.58 | 25.52–25.66 | 3.098 | 29.66× |
| scATAC Chromap | RapidMACS | 24 | 16.31 | 16.26–16.63 | 4.033 | 46.51× |
| Bulk ATAC ENCODE | MACS3 | 1 | 146.63 | 146.44–147.41 | 1.074 | 1.00× |
| Bulk ATAC ENCODE | RapidMACS | 1 | 41.94 | 41.80–42.00 | 1.755 | 3.50× |
| Bulk ATAC ENCODE | RapidMACS | 4 | 13.21 | 13.20–13.24 | 2.145 | 11.10× |
| Bulk ATAC ENCODE | RapidMACS | 24 | 7.47 | 7.33–7.55 | 2.646 | 19.63× |
| Bulk ATAC Chromap | MACS3 | 1 | 122.92 | 122.85–124.58 | 1.064 | 1.00× |
| Bulk ATAC Chromap | RapidMACS | 1 | 32.51 | 32.48–32.55 | 1.736 | 3.78× |
| Bulk ATAC Chromap | RapidMACS | 4 | 10.84 | 10.84–10.98 | 2.095 | 11.34× |
| Bulk ATAC Chromap | RapidMACS | 24 | 6.56 | 6.54–6.87 | 2.636 | 18.74× |
| CUT&RUN | MACS3 | 1 | 64.74 | 64.58–64.93 | 0.347 | 1.00× |
| CUT&RUN | RapidMACS | 1 | 9.94 | 9.93–9.94 | 0.458 | 6.51× |
| CUT&RUN | RapidMACS | 4 | 2.99 | 2.99–3.02 | 0.505 | 21.65× |
| CUT&RUN | RapidMACS | 24 | 2.01 | 2.01–2.06 | 0.610 | 32.21× |
| ChIP-seq | MACS3 | 1 | 616.90 | 616.79–623.18 | 2.745 | 1.00× |
| ChIP-seq | RapidMACS | 1 | 97.08 | 97.04–97.16 | 4.762 | 6.35× |
| ChIP-seq | RapidMACS | 4 | 27.38 | 27.25–27.43 | 4.906 | 22.53× |
| ChIP-seq | RapidMACS | 24 | 12.18 | 12.13–12.31 | 5.398 | 50.65× |

### S5 Stage-level timings

Table S4 breaks a RapidMACS run down by pipeline stage for the three configurations that exercise the distinct code paths: a streaming fragment input, and controlled BAMPE input with and without a matched control. The stages listed are the dominant ones rather than an exhaustive partition, so they need not sum to the total. Each figure is the time RapidMACS spends in that stage; it is not an attribution of the speedup to an individual change, which would require running the caller with each optimization disabled in turn. A dash marks a stage that does not arise separately on that path: on fragment input the treatment pileup and local background are built in the same sweep as the signal tracks and are counted on that row.

#### Raw data

All 72 measured invocations, per-run checksums, stage profiles, host metadata, and the exact benchmarked executable hash are archived with the release at https://github.com/morphic-bio/rapidmacs.

**Table S3.** Controlled assays measured against released MACS3 v3.0.3 and against the same version with the extra file removed. RapidMACS is unchanged between the two and is shown once. Medians of three warm-cache runs after one unrecorded warm-up, same host and NVMe placement throughout.

| Assay | MACS3 build | MACS3<br>Wall (s) | Speedup at $T =$ | | |
| --- | --- | --- | --- | --- | --- |
|  |  |  | 1 | 4 | 24 |
| ChIP-seq | as released | 616.90 | $6.35\times$ | $22.53\times$ | $50.65\times$ |
| ChIP-seq | extra file removed | 409.26 | $4.22\times$ | $14.95\times$ | $33.60\times$ |
| CUT&RUN | as released | 64.74 | $6.51\times$ | $21.65\times$ | $32.21\times$ |
| CUT&RUN | extra file removed | 60.65 | $6.10\times$ | $20.28\times$ | $30.17\times$ |

**Table S4.** Median RapidMACS stage times (s), same runs as Table S2.

| Pipeline stage | scATAC (FRAG) |  | CUT&RUN |  | ChIP-seq |  |
| --- | --- | --- | --- | --- | --- | --- |
| | $T=1$ | $T=24$ | $T=1$ | $T=24$ | $T=1$ | $T=24$ |
| Input parsing | 7.03 | 7.03 | 6.73 | 1.39 | 25.79 | 3.49 |
| Track construction | 28.92 | 3.28 | 0.06 | 0.06 | 0.05 | 0.06 |
| Treatment pileup | — | — | 1.66 | 0.25 | 1.65 | 0.23 |
| Local background $\lambda$ | — | — | 0.98 | 0.14 | 65.49 | 7.03 |
| $p$-scores and peak regions | 17.03 | 5.04 | 0.49 | 0.17 | 4.01 | 1.25 |
| **Complete program** | **59.81** | **16.31** | **9.94** | **2.01** | **97.08** | **12.18** |

